# Body dissatisfaction shapes time perception and interoception interplay

**DOI:** 10.1101/2024.10.21.619381

**Authors:** Anna Rusinova, Maria Volodina, Kristina Terenteva, Vladimir Kosonogov

**Affiliations:** Center for Bioelectric Interfaces, HSE University, Moscow, Russian Federation; Federal Center for Brain and Neurotechnologies of the Federal Medical and Biological Agency, Moscow, Russian Federation; Institute for Cognitive Neuroscience, HSE University, Moscow, Russian Federation

**Keywords:** Time Perception, Emotion, Interoception, Awareness, Heart Rate, Body Image

## Abstract

This study explores the relationship between emotions, physiological responses, interoception, and time perception. We also aimed to understand how body dissatisfaction influences this interplay. Participants (N = 40) watched videos designed to evoke positive, neutral, and negative emotions while their heart rate was monitored via electrocardiography.

Our findings indicate that video duration perception varied depending on emotional content. Negative and neutral videos were perceived as shorter, while positive videos were estimated more accurately. Negative emotions were linked to a decrease in heart rate, suggesting a “freeze response,” while positive and neutral content did not significantly affect heart rate.

A significant correlation between heart rate and time perception was observed in participants with a normal body image, indicating that heart rate might serve as an internal clock. However, this correlation was absent in those with body image dissatisfaction, implying a weakened link between physiological states and time perception. Additionally, a strong negative correlation between heart rate and interoceptive accuracy was found in participants with body image dissatisfaction.

The results suggest that body image dissatisfaction disrupts the integration of emotional and physiological signals, affecting how we perceive and understand our internal bodily sensations and emotions.

## Introduction

The perception of time is crucial to human existence and our daily actions (Buhusi & Meck, 2005). Our ability to understand and measure time is essential for planning, coordinating, and performing our activities. Without an adequate perception of time, we would struggle to organize our tasks, interact with others, and adapt to changes in our environment and navigate and function effectively in the world.

Furthermore, the distortions in time perception observed in psychiatric and neurological disorders such as Parkinson’s disease, depression, bipolar disorder, anxiety disorders and schizophrenia are still poorly understood (Teixeira et al., 2013). For example, individuals with depression often focus on negative past experiences and frequently report that time seems to pass slowly or even feels like it has stopped (Ren et al., 2023). Similarly, patients with Parkinson’s disease also tend to perceive time as moving more slowly. On the other hand, anxiety can cause an accelerated perception of time, particularly during periods of high stress and arousal (Holman et al., 2023). People with attention deficit hyperactivity disorder might feel that time is slipping away faster or slower than it actually is (Ptacek et al., 2019). Stanghellini et al. found that patients with schizophrenia may describe their perception of time as lacking continuity, with moments of feeling disconnected from one another (Stanghellini et al., 2016). This can manifest as a loss of the immediate flow of time, making events feel isolated and unrelated, which contributes to difficulties in organizing daily activities and maintaining social interactions. Thus, the study of time perception is not only fundamental to understanding human cognition, but also holds significant potential for practical applications that can positively impact individual and societal well-being and have practical significance for the diagnosis and treatment of various psychiatric and neurological disorders.

The question of how we perceive time intervals has long existed, and we are only beginning to uncover complex mechanisms underlying this process, despite numerous studies on the subject. There are several hypotheses regarding the physiological basis of time perception.

According to the internal clock model, scalar expectancy theory (Gibbon, 1977), and the attentional gate model (Block & Zakay, 1997), the basic element of each time estimation process is a pacemaker, producing pulses at a certain rate, which are accumulated and stored in working memory. Comparing the number of accumulated pulses with the number of reference durations from previous learning processes yields a duration judgment (Gibbon et al., 1984). It has been proposed that the perception of duration is intrinsically represented by groups of neurons spread across different brain areas (Karmarkar & Buonomano, 2007; Paton & Buonomano, 2018).

Craig’s embodiment concept suggests a direct link between time perception and bodily processes (A. D. B. Craig, 2009). It postulates that our sense of time is connected to visceral processes because they share common neural substrates, such as the insular, medial prefrontal, and orbitofrontal cortices (A. D. Craig, 2003). The heart is a strong candidate for the role of the internal clock because it produces a consistent rhythm. Its rhythmic activity not only regulates blood flow, but also sends continuous signals to the brain. These signals are processed by interoceptive centers, which help integrate bodily rhythms with our perception of time.

Research into the effect of heart rate on the perception of time has yielded mixed and sometimes contradictory results. While some studies found no impact of heart rate on time estimation, others discovered significant correlations. To begin with, Hawkes et al. (1961) found that rhythmic stimulation could change the subjective perception of time intervals (Hawkes et al., 1961). This study used various drugs to alter participants’ physiological parameters and found a positive correlation between heart rate and estimated time. Cahoon and colleagues also altered participants’ heart rate by threatening them with electric shocks, which were not actually inflicted on them (Cahoon, 1969). They concluded that participants responded to their own heartbeat when determining the subjective pace of time. Another study examined the effect of heart rate on time estimation in divers, cyclists, and recreational athletes, finding significant correlations between time estimates and heart rate (Jamin et al., 2004). It has also been hypothesized that the activity of the sympathetic and parasympathetic nervous systems may influence time perception. One study found that increased activity of the sympathetic system, indicated by higher heart rate and increased skin conductance response frequency, was associated with faster time perception (Ogden et al., 2022). However, some scientists question the relationship between time perception and heart rate (Ochberg et al., 1964). They suggest that heart rate may play a minor or insignificant role in time perception (Surwillo, 1982). Some studies have shown that time perception might not be related to physiological cues like the heartbeat but is instead determined by external sensory stimuli and cognitive processes (Fontes et al., 2016). For example, a study comparing smokers and nonsmokers found that, although their heart rate differed, their time estimation was the same (Carrasco et al., 1998).

Overall, the research suggests that the relationship between time perception and internal rhythmic cues is complex and multifaceted, requiring further analysis to fully understand the interaction between physiological factors and time perception.

There are many different factors and mechanisms that can affect the perception of time, emotional state being one of them. Emotional states influence our perception of time by adapting internal mechanisms in response to significant events (Droit-Volet & Gil, 2009). Emotions can make time seem to move faster or slower than it actually does, altering our internal clock and leading us to either overestimate or underestimate the duration of an event. In particular, negative emotions such as anger can lead to overestimation of time, while certain individual differences in emotionality reinforce this effect (Tipples, 2008). Thus, negative emotional stimuli can elongate the perceived duration of temporal stimuli, unconsciously speeding up the internal clock (Yamada & Kawabe, 2011). High arousal emotions like anger can enhance environmental sensitivity and alertness, which in turn affects time perception.

Interestingly, the relationship between time assessment, musical preparation, and the emotional content of musical stimuli (funny or sad songs) has been revealed (Panagiotidi & Samartzi, 2013). The results showed that musical training affects the perception of time, allowing people to estimate the length of time more accurately; while the emotional content of a song also affects the assessment of time, especially in the case of non-musicians who overestimated the duration of sad songs and underestimated the duration in the case of fun songs. There is also evidence that overestimation of time occurs at a moderate level of arousal for positive and negative valences, but at the same time underestimation also occurs in a state of high arousal with a negative valence (Van Volkinburg & Balsam, 2014). This suggests that emotional stimuli affect the processing of temporal information in qualitatively different ways at different stages of processing temporal information. Moreover, time underestimation also occurs when people are engaged in enjoyable activities such as watching entertaining videos or participating in fun events (Gonidis & Sharma, 2017). Authors found that Internet and Facebook related stimuli can distort time perception due to attention and arousal related mechanisms.

Our subjective well-being also strongly affects how we perceive time. Time seems to speed up during pleasurable activities but drags on during periods of boredom (Wittmann, 2009). Social isolation, and stress can slow the perception of time, while high levels of social satisfaction and low levels of stress can, conversely, speed up its passage (Droit-Volet et al., 2020; Martinelli et al., 2020). Thus, our sense of time results from a complex interaction between specific cognitive functions and our momentary emotional states.

After all, the way we perceive time can also be connected to our body image, as temporal processing and body schemas are essential for integrating our actions and perceptions (Prinz & Hommel, 2001). It is revealed that underestimations in time perception are connected to psychological conditions characterized by diminished processing of high salience stimuli from the body (Di Lernia et al., 2018) Additionally, time passes more slowly in situations where the body is less represented compared to medium and high levels of embodiment (Unruh et al., 2023). To investigate how the representation of one’s own body affects time perception, Unruh and colleagues conducted a groundbreaking experiment using virtual reality. Participants were randomly assigned to experience different levels of embodiment: without an avatar (low), with only hands (medium), or with a high-quality avatar representing their body (high). Furthermore, negative body image or dissatisfaction with one’s body can distort the perception of internal bodily signals (Todd et al., 2021). This, in turn, may impact the perception of time.

As for the theory that interoceptive signals influence time perception (A. D. B. Craig, 2009), a recent study investigating the interplay between emotions and time perception found that internal attentional focus amplifies the effect of emotions on video duration estimation (Pollatos et al., 2014). The study revealed that fear tends to elongate perceived time, especially with an interoceptive focus, whereas amusement shortens it. However, this study did not include physiological data or information about participants’ interoceptive abilities.

The aim of the present study was to test the hypothesis that the effect of emotion on time perception is mediated by interoceptive signals. We also sought to test the hypothesis that attentional focus, interoceptive accuracy and personal traits modulate this effect.

## Methods

### Participants

The sample consisted of 40 participants ranging in age from 18 to 37 years, with thirteen men and twenty seven women and the mean age of 22.4 (SD = 6.89). Inclusion criteria were the following: participants were selected from individuals aged 18 to 40 years, participants should be without diagnosed mental illnesses or brain disorders and not taking drugs that affect the central nervous system, such as antidepressants or sedatives. Exclusion criteria were the participant’s inability to complete the experiment and the incompatibility of data due to specific physiological parameters affecting data acquisition. Two participants were excluded from the analysis because their interoceptive accuracy scores failed to meet the criteria of the statistical test for outliers. Thus, the data from 38 individuals were included in the final analysis with twelve men and twenty six women and the mean age of 22.0 (SD = 6.58). The experiment was conducted in accordance with the Declaration of Helsinki and approved by the HSE University Committee on Inter-University Surveys and Ethical Assessment of Empirical Research (#52). Participants received compensation (8.33 USD at purchasing power parity). All participants provided written informed consent.

### Stimuli

In the main phase of the experiment, participants watched 36 video clips chosen from a pre-tested database. See Supplementary materials for descriptive data on the videos used in the current study).

Ratings of valence in that mentioned affective video database (from 1 for “very negative” to 9 for “very positive”) were M ± SD negative = 2.61 ± 0.16; M ± SD positive = 7.38 ± 0.21; M ± SD neutral = 4.89 ± 0.27.

Ratings of arousal (from 1 for “very calm” to 9 for “very arousing”) were M ± SD negative = 5.96 ± 0.25; M ± SD positive = 6.02 ± 0.37; M ± SD neutral = 3.81 ± 0.71. Negative and positive videos did not differ in arousal.

Negative videos were rated as more negative (t = -24.78, p < 0.001, d = -10.56) and arousing (t = 9.92, p < 0.001, d = 4.23) than the neutral ones. Positive videos were rated as more positive (t = 25.07, p < 0.001, d = 10.69) and arousing (t = 9.56, p < 0.001, d = 4.08) than the neutral ones.

These videos were categorized based on their emotional impact: 12 evoked positive emotions, others 12 evoked negative emotions, and the third group of 12 was neutral, as determined by prior ratings. For each category, we selected two videos for the following durations: 21, 28, 35, 42, 49, and 56 seconds.

### Procedure

The design of the experiment is shown in Figure 1.

**Figure 1.**
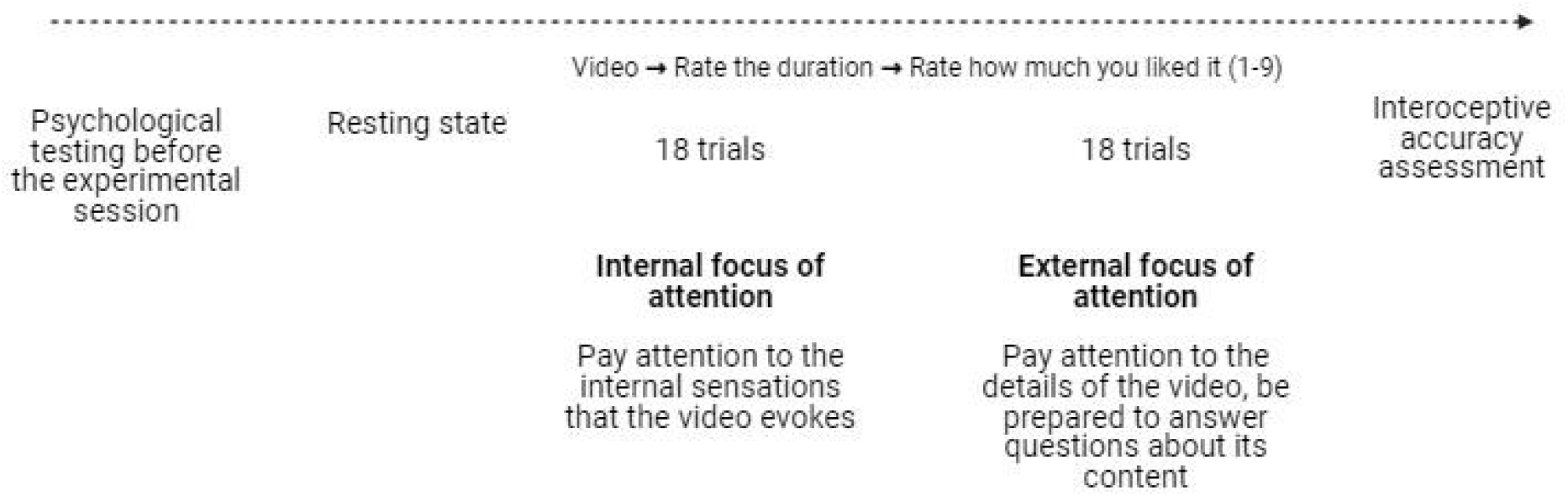
The design of the experiment. The sequence of internal and external attention focus varied among participants.

Before the experimental session, all participants completed a series of psychological questionnaires (see below). Then, participants were given three minutes to rest and minimize any activity. This resting phase aimed to stabilize their physiological performance before they were exposed to the experimental stimuli, allowing for a more accurate investigation of how visual content affects cognitive processes.

The experimental session was divided into two equal parts. In the first part, participants focused on the internal sensations triggered by the videos (internal attention focus). In the second part, they concentrated on the video’s details (external attention focus). The order of these conditions varied among participants.

After watching each video, participants rated its duration in seconds and assessed their feelings about the content on a scale from 1 to 9, where 1 represented “very unpleasant” and 9 represented “very pleasant.”

The final phase of the study involved a task designed to evaluate interoceptive accuracy. Participants were instructed to concentrate on their own heart rate and count their heartbeats over specific durations of 21, 28, 35, 42, 49, and 56 seconds (Garfinkel et al., 2015). Interoceptive accuracy was evaluated according to the following formula:

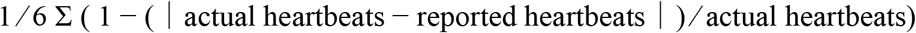

Presentation of the stimulus material and recording of the subjects’ responses were performed using the Psychopy program (Peirce, 2007). ECG was continuously monitored throughout the experimental session.

### ECG recording

Throughout the experiment, ECG was recorded using an NVX52 amplifier (Medical Computer Systems, Russia) and NeoRec software (Medical Computer Systems, Russia) and digitized at a sampling rate of 250 Hz. Clamp ECG electrodes were positioned on the wrists and the ankle of the right foot. Synchronization of the ECG signal and stimulus presentation was performed using a photo sensor fixed in the upper right corner of the monitor.

### Psychological questionnaires

Before the experimental session, all participants completed a series of psychological questionnaires designed to assess level of anxiety, empathy, body awareness, and body image. These factors can influence the perception of interoceptive signals and the response to emotionally charged stimuli. Full text of the questionnaires can be found in Supplementary materials.

#### The Questionnaire Measure of Emotional Empathy

(QMEE) by Mehrabian and Epstein (Mehrabian & Epstein, 1972). The participants completed the Russian version by Stoliarenko (Kosonogov, 2014). The questionnaire is widely used in psychology and sociology. It shows the general capacity for empathy, the level of expression of reaction and the direction of experience. The Russian QMEE consists of 25 items designed to gauge an individual’s tendency to respond emotionally to the experiences of others in various situations (e.g., “Sometimes the words of a love song can move me deeply”).

#### Multilevel Assessment of Interoceptive Awareness, MAIA-R

The Russian adaptation of the MAIA-R Multilevel Assessment of Interoceptive Awareness MAIA-R (Mehling et al., 2012) by Popova and Lopukhova (Popova & Lopukhova, 2022) was used to measure interoceptive awareness. The 32-item instrument uses a 6-level Likert scale response format (0 = never, 5 = always;). In the Russian version, Cronbach’s alpha reliability coefficient, calculated for each of the scales, ranges from 0.58 to 0.85, and for the entire questionnaire it is 0.89, indicating a high overall consistency of the questionnaire.

#### The Body Image Questionnaire

by Skugarevsky and Sivukha was used to assess the level of dissatisfaction with one’s own body (Skugarevsky & Sivukha, 2006). Respondents are asked to rate 16 items on a 4-point scale (from 0 for “never” to 3 for “always”). The internal consistency of the items in our sample is excellent, α = 0.95. In the article dedicated to the development of this tool, the authors provide a threshold value of 13 points, which may indicate disordered eating behavior and is recommended for use in screening studies, as well as an evaluation of the risk at 32 points, which indicates a high level of body dissatisfaction and a high risk of eating disorder in the respondent.

#### The Hamilton Anxiety Rating Scale (HAMA)

quantifies the severity of anxiety and is often used to evaluate antipsychotic medications (Thompson, 2015). It consists of 14 indicators, each of which is defined by a number of symptoms. Each indicator is rated on a 5-point scale from 0 (absent) to 4 (severe).

#### Mindful Attention Awareness Scale (MAAS)

A Russian adaptation of Mindful Attention Awareness Scale (Brown & Ryan, 2003) by Golubyev (Golubev, 2012) was performed to measure mindfulness. MAAS is a 15-item self-report tool designed to measure mindfulness, which is defined as a process of increased receptive awareness and attention to the present moment. Each item is rated on a six-point Likert scale from 1 (“almost always”) to 6 (“almost never”) and then added together to create a total sum score. Thus, lower scores indicate greater mindlessness, whereas higher scores indicate greater mindfulness.

#### Emotionality questionnaire

This diagnostic of emotionality was proposed by Suvorova in 1976 and determines the general emotionality of a person (Suvorova, Valentina Vasilyevna-Tests for determining individual characteristics of vegetative response). The methodology includes 15 questions (statements). A score of 1 is assigned to each affirmative answer. The total sum of points is calculated.

#### Beck’s Depression Inventory

This instrument is a 21-item, self-report questionnaire designed to assess and evaluate the frequency of anxiety symptoms over a one-week period (Beck et al., 1988). This test assesses two factors: cognitive and somatic symptoms. The instrument has good internal consistency (α = 0.92), test-retest reliability (r = 0.75; df = 81, P = <.001), and convergent and discriminant validity.

### ECG data processing

The following filters were used: 0.5 Hz high-pass filter, 25 Hz low-pass Butterworth filter, notch filter at 50 Hz. R-peaks on the ECG were detected using the bio_process function from the NeuroKit library in Python (Makowski et al., 2021).

### Data analysis

Statistical analysis was conducted using Python scripts and the Statistica 64 software. Initially, we examined the relationships between video pleasantness and heart rate, video pleasantness and duration estimation error, HR and duration estimation error, focus of attention and HR, and focus of attention and duration estimation error across the entire sample in the study. This analysis utilized mixed linear model regression, incorporating subject ID as a random factor to account for individual differences. The statsmodels.formula.api.mixedlm function from Python was used.

Then, based on subjective ratings of video pleasantness, the videos were categorized as negative (scores less than 4 out of 9), neutral (scores ranging from 4 to 6), and positive (scores between 7 and 9). A repeated measures analysis of variance (ANOVA) was conducted to assess the effect of video valence on duration estimation error and HR.

The correlation between the averaged across all participants normalized (z-score) HR and the pleasantness rating of the videos, as well as the correlation between a person’s interoceptive accuracy and their average duration estimation error for the stimulus, and person’s interoceptive accuracy and personal traits were assessed using Pearson correlation analysis.

To identify whether personality traits influence a person’s reaction to the shift in focus from external to internal attention, we calculated the average difference in HR between the two conditions for each individual. We then conducted a correlation analysis to determine the relationships between this difference and the scores of psychological questionnaires.

In the second part of the study, the analysis described above was conducted separately for two subgroups of participants: those with body image scores within the normal range (less than 13) and those with significant body dissatisfaction as indicated by the questionnaire.

## Results

### Video stimuli pleasantness and time perception

Initially, we explored how the pleasantness of the video impacted the participants’ perception of its duration. To achieve this, we examined the relationship between participants’ subjective ratings of video pleasantness and their accuracy in estimating video duration. A negative error indicated an underestimation of the video’s length, while a positive error reflected an overestimation.

The mixed linear model regression analysis with a random factor revealed that the subjective evaluation of pleasantness of video was significantly associated with estimation error of video duration, β = 0.013, SE = 0.000, z-score = 5.140, p < 0.001, 97.5% CI [0.008, 0.018]. According to subjective ratings of video pleasantness, videos were labeled as negative (score less than 4 out of 9), neutral (4 to 6 points), and positive (7 to 9 points). Repeated measures ANOVA analysis revealed a significant effect of video valence on the estimation error of video duration, F(2.74) = 9.205, p = 0.0003. Participants underestimated the durations of negative and neutral videos. The absolute estimation error of negative (Tukey test, p = 0.0003) and neutral (Tukey test, p = 0.04) videos were significantly larger than the estimation error of positive videos (Figure 2). Meanwhile, no interaction between the factors “video valence” and “focus of attention” was found.

**Figure 2.**
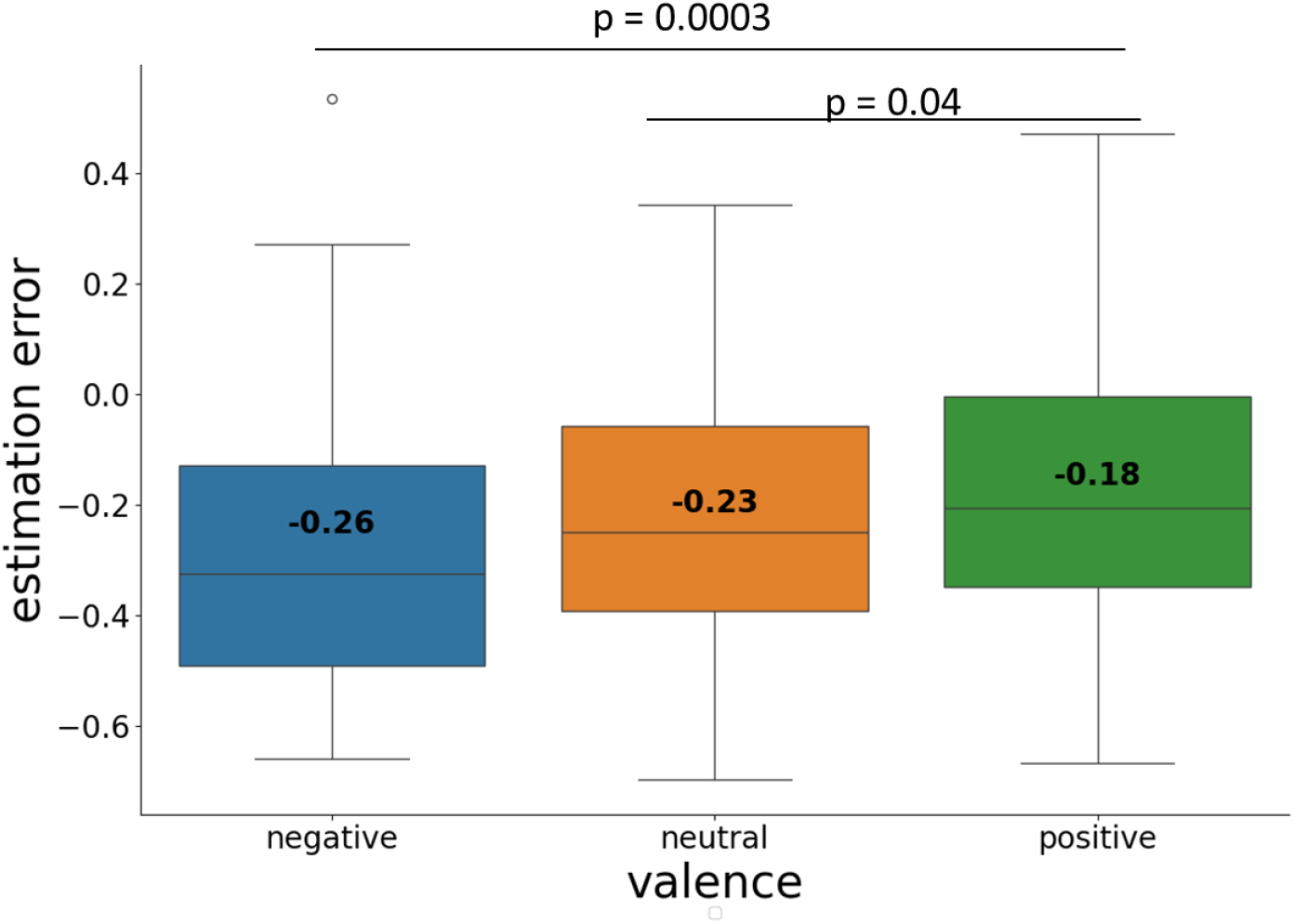
Error of video duration estimation when viewing negative, neutral, and positive videos. The box indicates the interquartile range (IQR), with the lower edge representing the 1st quartile (Q1) and the upper edge representing the 3rd quartile (Q3). The box contains the middle 50% of the data and medians, while the whiskers extend to the minimum and maximum values. The indicated numbers represent the mean values for each group. The p-values for the post-hoc Tukey test are provided.

### Video stimuli pleasantness and heart rate

Next, we examined the relationship between the perceived pleasantness of the video and the participants’ heart rate during its viewing. The pleasantness factor was significantly associated with normalized heart rate (z-score) (β = 0.043, SE = 0.014, z_score = 3.146, p = 0.002, 97.5% CI [0.016, 0.069]), with participant ID included as a random effect.

Repeated measures ANOVA analysis indicated a significant effect of video valence on the normalized HR, F(2.74) = 5.7, p = 0.005. Heart rate decreased while watching videos eliciting negative emotions compared to eliciting neutral (Tukey test, p = 0.01) and positive emotions (Tukey test, p = 0.01, Figure 3).

**Figure 3.**
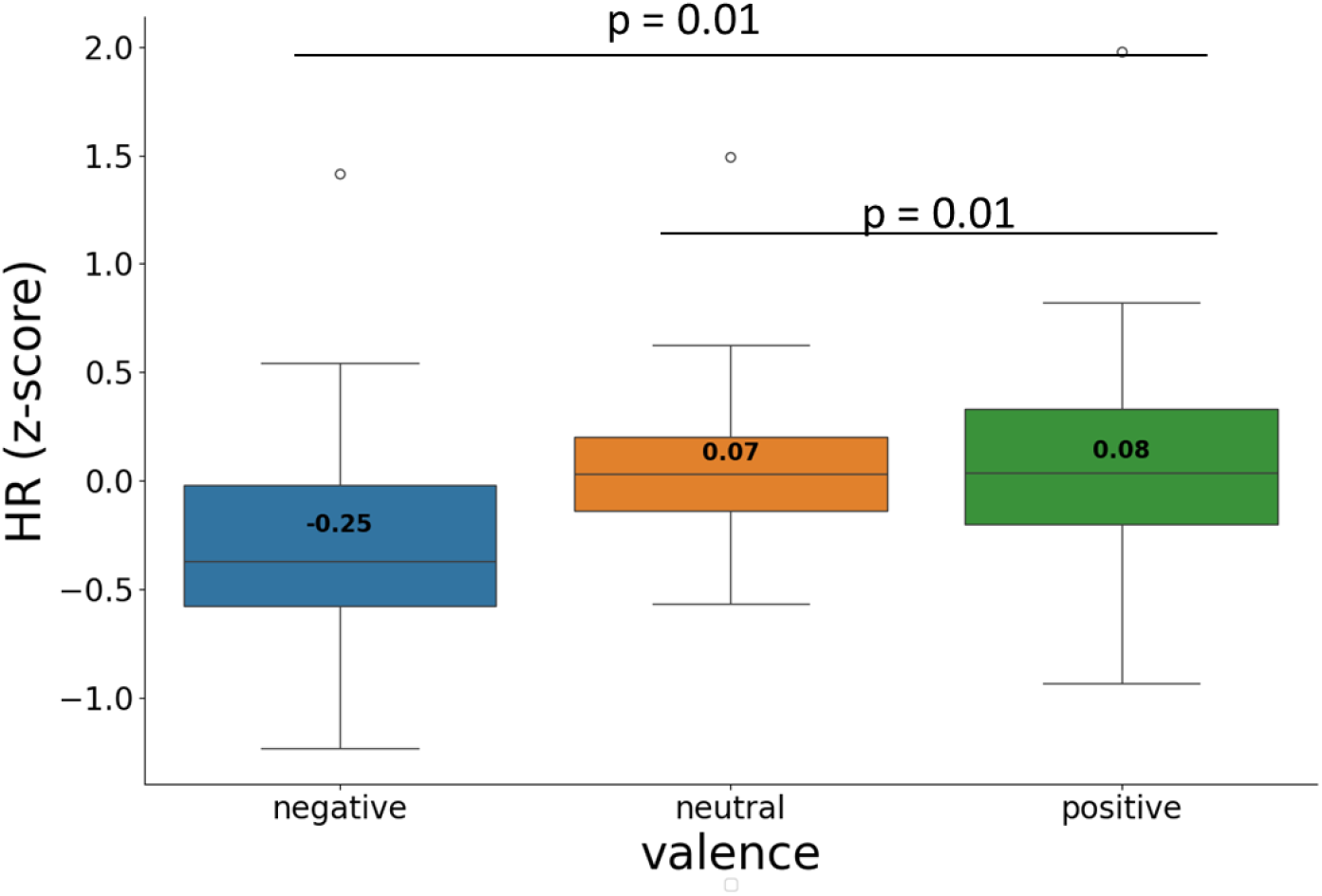
Error of video duration estimation when viewing negative, neutral, and positive videos. The box indicates the interquartile range (IQR), with the lower edge representing the 1st quartile (Q1) and the upper edge representing the 3rd quartile (Q3). The box contains the middle 50% of the data and medians, while the whiskers extend to the minimum and maximum values. The indicated numbers represent the mean values for each group. The p-values for the post-hoc Tukey test are provided.

This is consistent with the results of the correlation analysis, which demonstrated a direct correlation between the subjective evaluation of video pleasantness averaged across all participants and the normalized HR observed while watching a given video, r = 0.37, p = 0.028. Thus, videos rated as unpleasant caused a slowing of the heart rate, while videos rated as pleasant, on the contrary, accelerated the HR (Figure 4).

**Figure 4.**
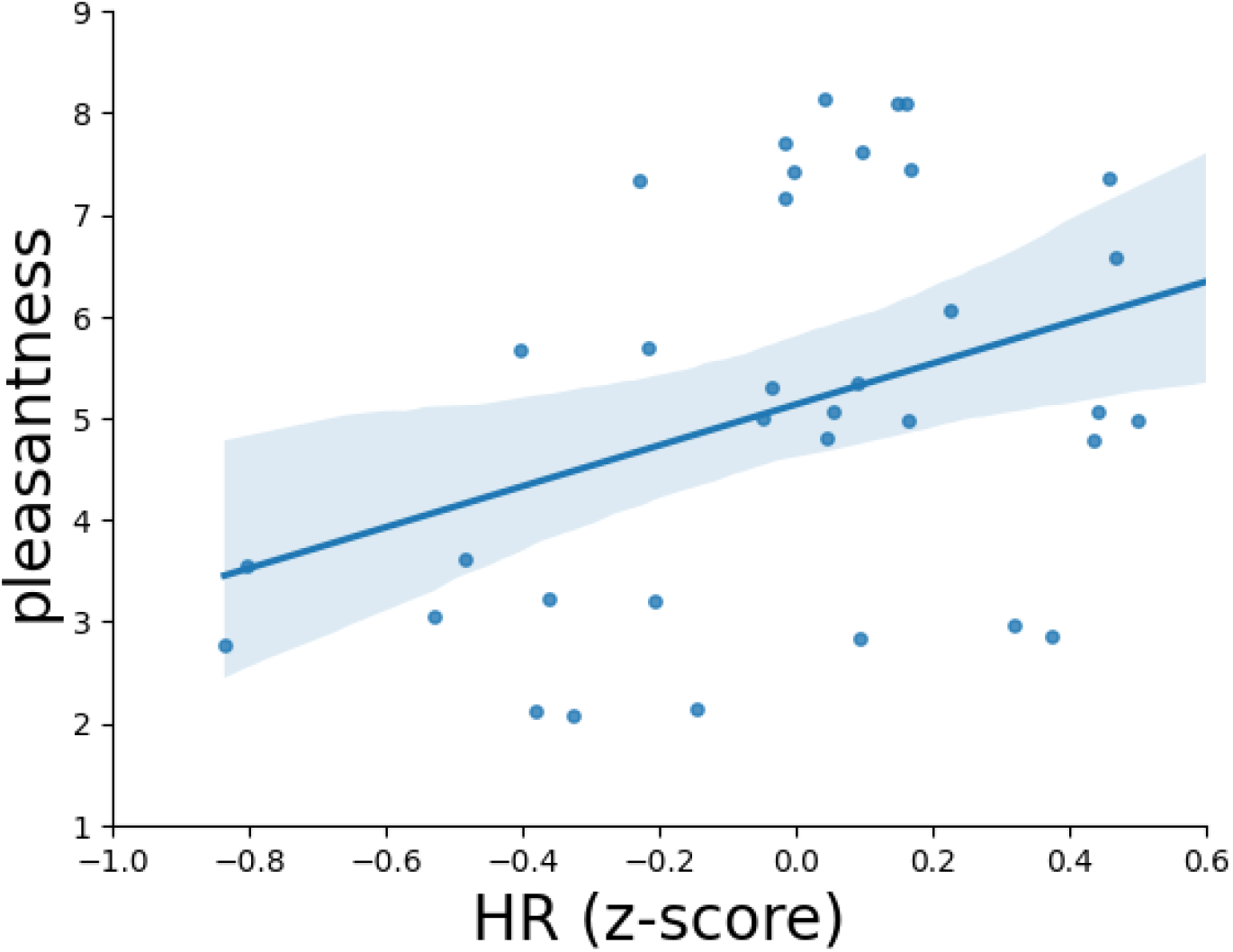
Correlation between the mean normalized heart rate of all subjects and the mean pleasantness scores of the video, r = 0.37, p= 0.014. One point corresponds to one video stimulus.

### Heart rate and time perception

In the next step, we examined the hypothesis concerning the relationship between heart rate and time perception. The mixed linear model regression analysis found that the normalized HR (z-score) factor was significantly associated with estimation error of video duration (β = 0.021, SE = 0.006, z score = 3.592, p < 0.001, 97.5% CI [0.01, 0.033]). Thus, the slower a person’s heart rate was, the less time they feel has passed.

When the data of only neutral videos were included in the model, the same pattern was observed, β = 0.021, SE = 0.009, z-score = 2.448, p < 0.014, 97.5% CI [0.001, 0.038], indicating that the effect of heart rate on time perception is independent or not fully dependent on emotional state.

### Interoceptive accuracy, time perception and personal traits

Other evidence in favor of the hypothesis that conscious perception of heartbeat can influence time perception would be the correlation of interoceptive accuracy and video duration estimation error. However, correlation analysis revealed no significant correlation between participants’ interoceptive accuracy and the participant’s average video duration error recorded during the experiment (all ps > 0.05).

Moreover, the correlation analysis performed on the entire sample failed to identify any meaningful relationships between interoceptive accuracy, errors in estimating stimulus duration, and scores from the psychological questionnaires (all ps > 0.05).

### Focus of attention, estimation error, and HR

The mixed linear model regression analysis with a random factor found that internal attention focus was associated with decreased estimation error of video duration (β = - 0.026, SE = 0.012, z score = - 2.236, p = 0.025, 97.5% CI [-0.049, -0.003]) and decreased normalized heart rate (β = - 0.176, SE = 0.056, z score = - 3.151, p = 0.002, 97.5% CI [-0.286, -0.067]). However, there was no significant interaction between focus direction, normalized HR and pleasantness factors. Attention direction did not modulate the interplay between heart rate and time perception.

### A person’s perception of their body matters

We posited that the effects of attention focus could be linked to specific psychological traits. To test our hypothesis regarding normalized heart rate and duration estimation error, we calculated the differences between the external and internal focus of attention across various conditions (during watching of negative, neutral and positive stimuli). Next, using correlation analysis, we assessed which psychological measures correlate with the obtained values. We found that body image questionnaire scores were positively correlated with the difference in normalized heart rate between external and internal focus of attention conditions, r = 0.43, p = 0.007 (0.09 after FDR correction; Figure 5). High scores in the body image questionnaire correspond to a state of dissatisfaction with one’s own body.

**Figure 5.**
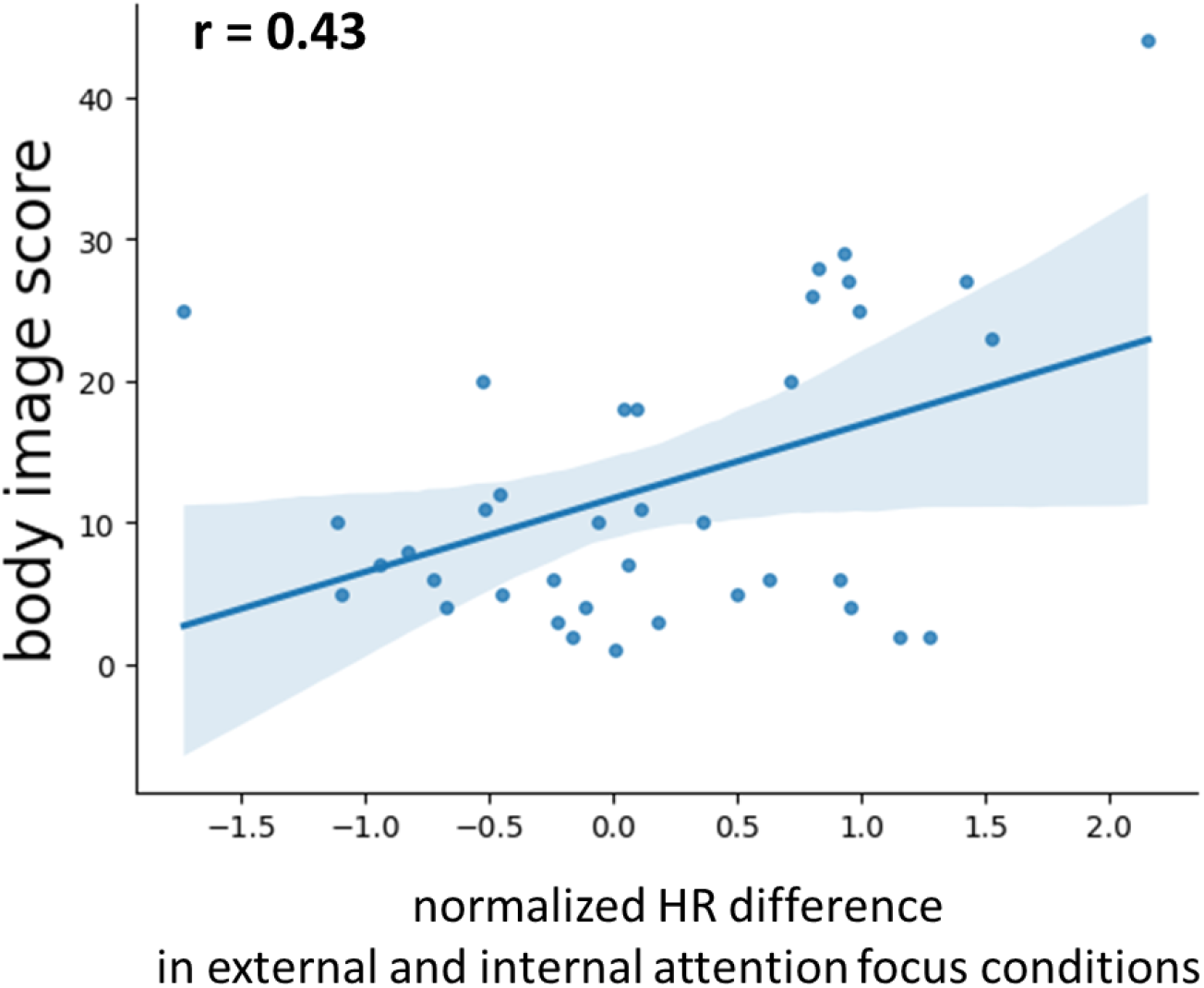
Correlation of normalized HR difference in external and internal attention focus conditions and body image questionnaire scores.

Our results revealed that people with impaired body image (questionnaire score > 12) have a pronounced difference in heart rate under conditions of attention directed to an external stimulus and to their internal sensations. This effect was more pronounced under emotional load. In the case of inward attention, they showed a decrease in HR, as well as all participants did when watching videos that evoke negative emotions (Figure 6).

**Figure 6.**
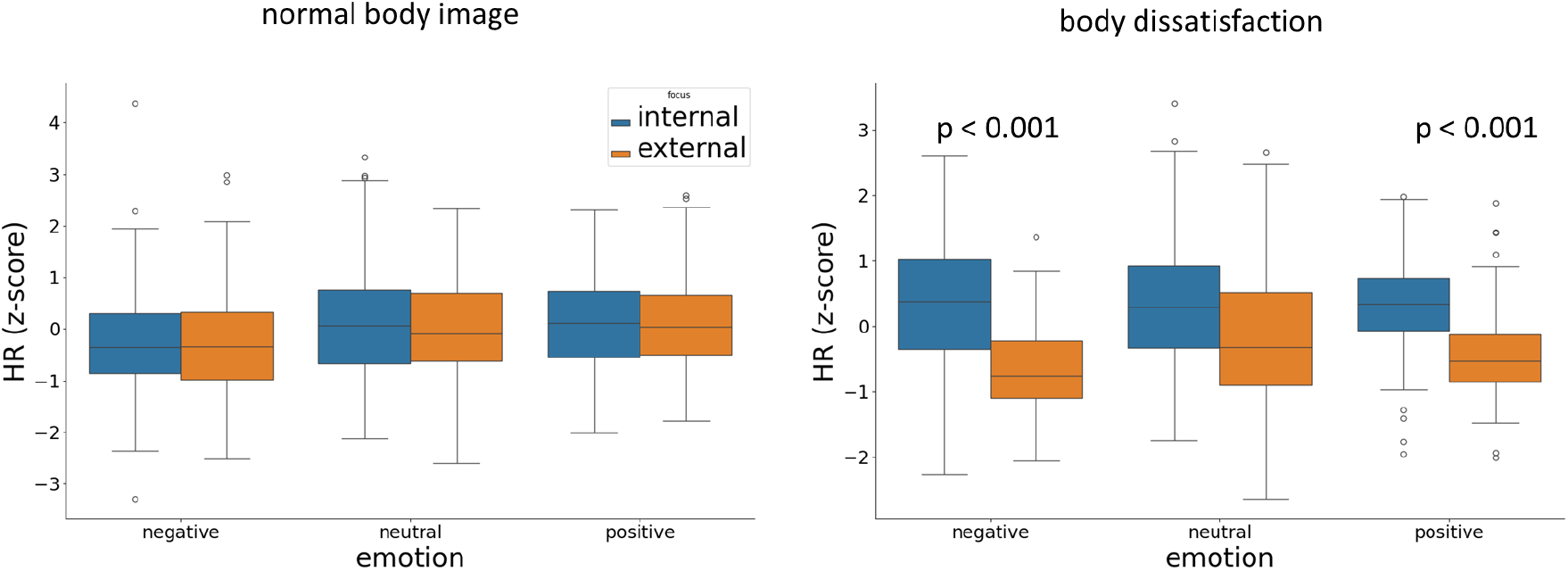
Normalized HR under internal and external attention conditions during watching videos in people with normal body image (questionnaire score < 13, n = 25) and people with body dissatisfaction (questionnaire score 13 or more, n = 13). The box indicates the interquartile range (IQR), with the lower edge representing the 1st quartile (Q1) and the upper edge representing the 3rd quartile (Q3). The box contains the middle 50% of the data, while the “whiskers” extend to the minimum and maximum values. The indicated numbers represent the mean values for each group. The p-values for the post-hoc Tukey test are provided and correspond to differences between internal and external attentional focus.

In light of the differences observed between subgroups with a normal body image and those experiencing body dissatisfaction, we conducted a separate reanalysis of each subgroup.

Body image dissatisfaction did not significantly influence the relationship between video pleasantness and time perception, nor between video pleasantness and heart rate (all ps > 0.05). However, while analyzing the correlations between heart rate and duration estimation error, as well as heart rate and interoceptive accuracy, we observed intriguing patterns specifically within the subgroup of participants with high body image scores.

Correlation analysis indicated a positive correlation between persons’ heart rate averaged over all epochs and average time estimation error in the subgroup of individuals with a normal body image. This finding supports the original hypothesis that the heartbeat serves as an internal metronome: as heartbeats slow, subjective time perception also slows. In contrast, no association was found between heart rate and estimation error in the subgroup with body image dissatisfaction (Figure 7).

**Figure 7.**
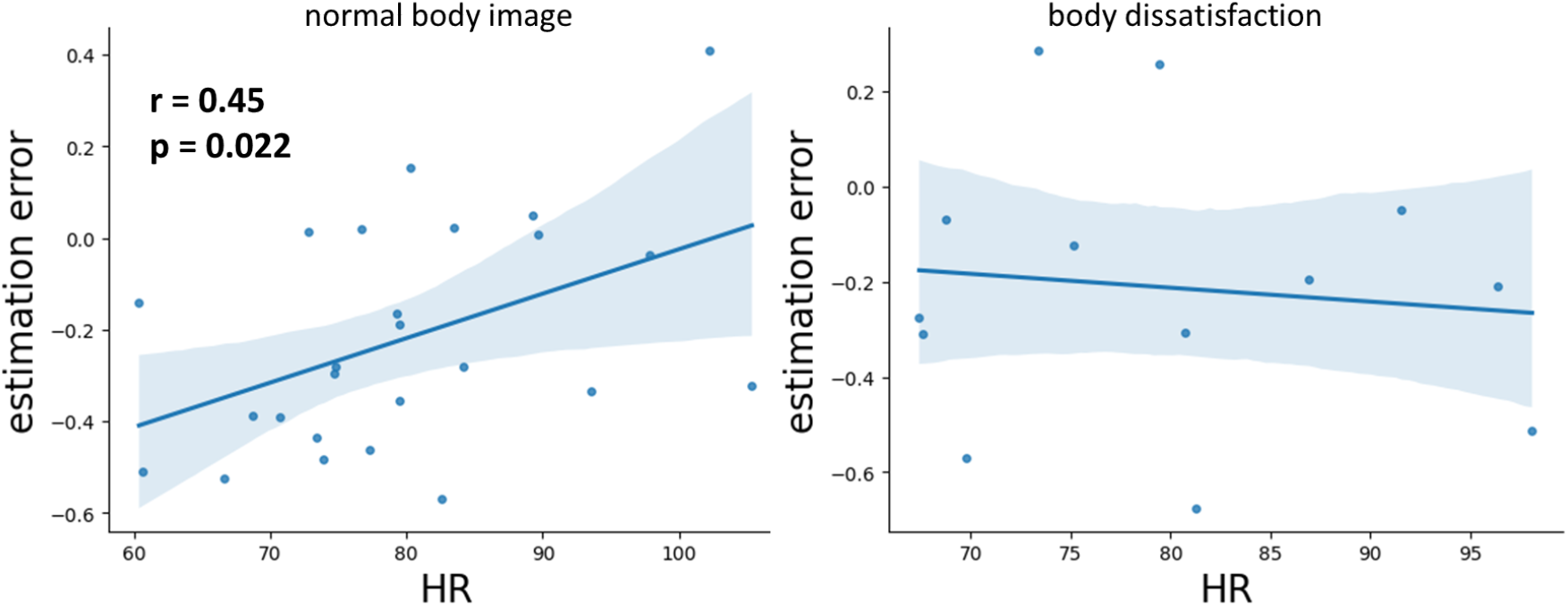
Correlation of mean HR and mean estimation error averaged across all epochs in participants with normal body image questionnaire score (left, n = 25) and high body image scores (right, n = 13). One point corresponds to one subject.

Another intriguing finding was that correlation analysis revealed a strong negative correlation between mean heart rate (HR) and interoceptive accuracy in the subgroup of individuals who scored high on the body image questionnaire. In contrast, no correlation was observed between these two measures in the group with a normal body image (Figure 8).

**Figure 8.**
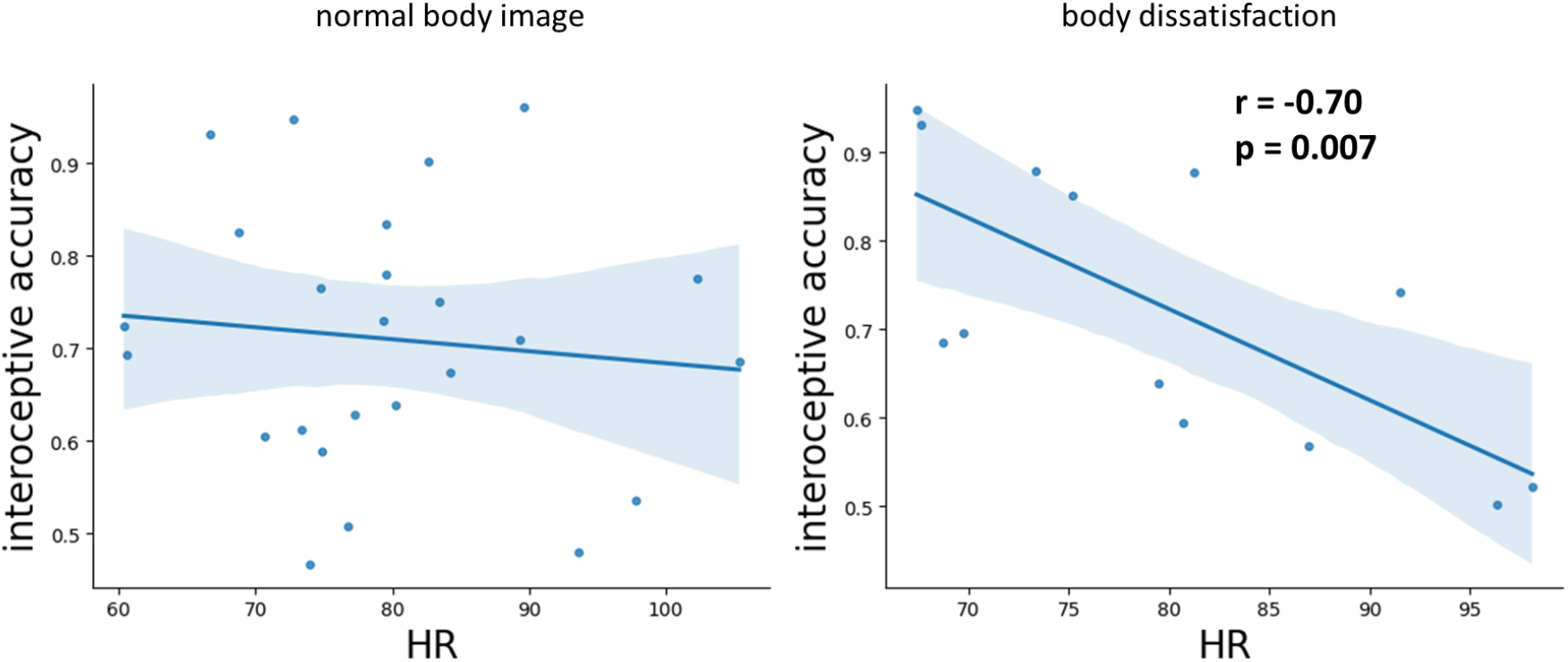
Correlation of HR averaged across all epochs and interoceptive accuracy in participants with normal body image questionnaire score (left, n = 25) and high body image scores (right, n = 13). One point corresponds to one subject.

## Discussion

The goal of this study was to explore the hypothesis that interoceptive signals mediate the relationship between emotion and time perception. We also aimed to investigate how factors such as attentional focus, interoceptive accuracy, and individual personality traits may influence this relationship.

Our results confirmed that the perception of video duration can vary greatly depending on emotional content. Negative and neutral videos seemed shorter to participants than they actually were, while the estimation errors for positive videos were smaller. The impact of emotional state on time perception has been previously documented in several studies (Tipples, 2008; Yamada & Kawabe, 2011; Droit-Volet & Gil, 2009).

However, the direction of this effect has varied among these studies. In some cases, similarly to ours, negative emotions resulted in an underestimation of the time intervals. Thus, in a study by Sarigiannidis et al. (Sarigiannidis, Grillon, et al., 2020), induced anxiety led to underestimation of duration, thus, time seemed to pass quicker. Similarly, studies by Tamm et al. (Tamm et al., 2014) and Van Volkinburg et al. (Van Volkinburg & Balsam, 2014) also found that negative emotions result in the underestimation of time intervals.

However, some studies reported an opposite effect. Anger and fear led to overestimation of the perception of time, and this effect was influenced by individual differences in negative emotionality (Tipples, 2008; Pollatos et al., 2014). Also, it was found that anxious people feel that time passes more slowly when they are exposed to short presentations of threatening stimuli (Bar-Haim et al., 2010).

This discrepancy may stem from the various types and durations of stimuli presented, along with the different emotional nuances experienced by participants in the studies.

However, it is important to recognize that different motivational tendencies within negative states should be studied meticulously in the future. Experiments show that negative affect and motivational directionality affect time perception differently (Gable et al., 2016). Based on the motivational direction model of time perception, it was proposed that sadness and anger, hypothesized approach-motivated negative states, cause time to shorten, but disgust, a withdrawal-motivated negative state would cause time to lengthen. The similar effect of sadness (Benau & Atchley, 2020) and depression scores (Gil & Droit-Volet, 2009) was also found.

One hypothesis for the underestimation of time experienced during negative emotions is that individuals may misjudge the passage of time when their attention is focused inward, concentrating on their internal experiences caused by negative stimuli. Our findings support this hypothesis, demonstrating that an internal focus of attention is linked to an underestimation of time duration. There is an assumption that time passes more quickly by overloading neural resources, particularly in the mid-cingulate cortex, potentially driving emotion-related changes in temporal perception (Sarigiannidis, Kieslich, et al., 2020).

Specifically, when a person experiences a strong emotion, their brain may become overloaded with processing that emotion, causing time to seem to pass faster.

Another hypothesis suggests that the physiological effects of emotions, such as fluctuations in heart rate, may also impact time perception.

It is widely recognized that individuals experiencing negative emotions frequently exhibit an elevated heart rate (Wu et al., 2019; Karki & Mahara, 2022). However, our findings reveal a different phenomenon: watching videos that evoke negative emotions was associated with a decrease in heart rate, in contrast to videos featuring positive or neutral content. This phenomenon may be linked to the “freeze response,” a physiological reaction characterized by heightened attention and concentration. During this response, a decrease in heart rate can enhance our ability to detect and respond to potential threats, serving as a vital component of our stress-response system. This adaptive mechanism allows individuals to remain alert in potentially dangerous situations (Kim et al., 2018). Empirical evidence supports the notion that a slow heart rate can facilitate a heightened state of awareness. For instance, individuals displayed a freeze reaction alongside a decline in heart rate in response not only to physical threats but also in response to social threats, like angry faces (Roelofs et al., 2010; Noordewier et al., 2020).

Additionally, responses to negative emotional stimuli often involve increased parasympathetic activity, which may lead to cardiac deceleration (Hagenaars et al., 2014). Research has shown that emotions such as disgust and sadness can activate the amygdala and are associated with this physiological response (Rohrmann et al., 2009). In certain circumstances, a reflex known as the vasovagal response may occur, characterized by simultaneous reductions in both heart rate and blood pressure in response to negative emotions (Dani et al., 2021). A similar decrease in heart rate in response to negative emotions was found in mentally retarded children (Corona et al., 1998). Such reactions can manifest in circumstances where an individual attempts to dissociate from their environment. When faced with some stressors, individuals may enter a state of “freezing,” which serves to minimize their visibility to potential dangers and facilitates a temporary emotional disengagement. In future research, it would be highly beneficial to gather more detailed information from participants regarding the emotions they experience while watching videos. This information would be useful for drawing conclusions about the influence of specific types of positive and negative emotions on both physiological reactions and time perception.

In our study, we identified a significant association between heart rate and participants’ perception of time intervals. Our results showed that as heart rate decreased, participants tended to underestimate the duration of time intervals more significantly. This finding is consistent with a body of research conducted previously (Hawkes et al., 1961). Thus, our finding supports the hypothesis that heartbeats serve as a kind of time-keeper, influencing the subjective experience of time. Moment-to-moment experience of time is synchronized with, and changes with, the length of a heartbeat.

Despite evidence suggesting that interoceptive signals play a role in time perception based on data regarding the relationship between heart rate and perception of time, our analysis of the entire sample revealed no correlation between participants’ interoceptive accuracy and their mean duration estimation error. This contrasts with previous studies that identified such a relationship (Uraguchi et al., 2022; Teghil et al., 2020).

An inward focus of attention was associated with a more pronounced underestimation of video duration and a lower normalized heart rate. However, we did not observe any significant interactions between focus direction, normalized heart rate, and factors related to pleasantness, unlike studies by Pollatos et al. (Pollatos et al., 2014) and Tamm et al. (Tamm et al., 2014), which indicated that an internal focus of attention amplifies the influence of emotional state on duration estimation error.

Based on our findings, it can be assumed that the absence of a modulatory effect of attentional focus on the relationship between emotions and timing errors is related to the fact that the influence of interoceptive signals on time perception operates at a subconscious level, whereas interoceptive awareness contributes little to this process.

The role of internal attention focus in time perception was also highlighted in a recent study (Kramer et al., 2013). The findings indicate that enhanced attention to internal states during meditation practice can lead to a relative overestimation of time duration, suggesting that focusing on internal sensations does not enhance accuracy in time estimation. Similarly, our results demonstrate that an internal focus of attention disrupts the ability to accurately estimate time duration; however, the direction of the error in our study was different.

After failing to identify the anticipated modulatory effect of attentional focus on the relationship between emotions, time perception, and heart rate, we decided to explore whether specific psychological characteristics might be related to a more pronounced influence of attentional focus on physiological measures. By analyzing the correlation between the differences in physiological indicators under external and internal attention focus and the scores from self-report questionnaires, we discovered a significant relationship between the effect of attention focus and body image scores. Individuals with high body image scores, indicating notable dissatisfaction with their bodies, exhibited a greater effect of inward attention focus compared to those with normal scores on this measure. Internal attention can increase body awareness, which for people with a disturbed body image can be unpleasant and stressful, and this, in turn, leads to physiological changes. Thus, individuals with disrupted body image exhibited a freezing response during inward attention focus, similar to the reaction elicited by viewing unpleasant videos. This effect was more pronounced under emotional load.

Considering the differences between individuals with elevated scores and those with scores within the normal range, we conducted a separate analysis for the two groups of participants.

We conducted a correlation analysis to examine the relationship between persons’ mean heart rate and mean duration estimation error in two separate subgroups. Our findings revealed a significant correlation only in the subgroup of individuals with body image scores within the normal range. In contrast, among those with pronounced dissatisfaction with their bodies, heart rate did not correlate with time perception.

Therefore, while in individuals with a normal body image, the heart rate can act as an internal timekeeper, people with distorted body image lack this connection. Their internal feelings and physiological processes do not influence their perception of time. This may be because they have reduced interoception abilities and may avoid focusing on their body sensations due to negative thoughts about their body. A recent study found that negative body image disrupts interoception at an unconscious level (Todd et al., 2021). Weaker brain responses to the gut and heart were associated with greater levels of body shame and weight preoccupation, so people are less sensitive to their body sensations and focus more on the external perception of their body. Moreover, greater fear of bodily sensations also can lower the effect of self-awareness (Petersen & Ritz, 2011).

An additional distinction between individuals with body image dissatisfaction and those with normal body image scores was the strong negative correlation between mean heart rate and interoceptive accuracy. This correlation was found only in individuals with body image dissatisfaction.

This result can be explained by the fact that a high heart rate may be associated with increased levels of stress and anxiety (Trotman et al., 2019), this, in turn, can impair a person’s ability to accurately perceive their internal body signals (Back & Bertsch, 2020). Research has shown that individuals with eating disorders tend to experience a decrease in interoceptive awareness, potentially stemming from a negative perception of their bodies and a tendency to avoid bodily sensations (Datta & Lock, 2023). Furthermore, those with a heightened negative perception of their bodies may experience an underlying sense of anxiety or other negative emotional states, a phenomenon not typically seen in individuals with a normal body image (Todd et al., 2021).

In conclusion, our research on the relationship between emotions, heart rate, and time perception has revealed several key aspects. We established that heart rate and emotional states are closely linked to how individuals perceive time. Interoceptive signals originating from the heart play a significant role in this assessment, influencing how emotions shape our perception of time. Furthermore, our results indicate that attentional focus does not affect the relationship between emotions and time perception, suggesting that these influences may be tied to subconscious interoceptive signals rather than conscious interoception.

We also found that individuals with body perception disorders exhibit a disruption in this relationship, indicating that their ability to integrate emotional and physiological signals may be impaired. These disruptions could diminish their interoceptive processing, which in turn affects emotional regulation and time perception.

Ultimately, our findings underscore the complexity of the relationships among heart rate, emotions, interoceptive accuracy, and time perception. The role of unconscious interoceptive signals suggests that physiological states can influence us without our full awareness. Understanding these interconnections is crucial for developing effective strategies to address issues related to body perception and emotional well-being, paving the way for more integrated and accurate self-representations and perceptions of time.

## Supporting information

Questionnaires

## Acknowledgments

The authors would like to thank the participants for taking part in this research.

## Data availability statement

The data that support the findings of this study are available from the corresponding author upon reasonable request.

## Author contributions

Maria Volodina, Vladimir Kosonogov: Contributed to conception and design. Anna Rusinova, Kristina Terenteva: Contributed to acquisition of data. Maria Volodina, Anna Rusinova: Contributed to interpretation of data. Anna Rusinova, Maria Volodina, Kristina Terenteva, Vladimir Kosonogov: Writing— original draft, review and editing.

